# Engineered SARS-CoV-2 receptor binding domain improves immunogenicity in mice and elicits protective immunity in hamsters

**DOI:** 10.1101/2021.03.03.433558

**Authors:** Neil C. Dalvie, Sergio A. Rodriguez-Aponte, Brittany L. Hartwell, Lisa H. Tostanoski, Andrew M. Biedermann, Laura E. Crowell, Kawaljit Kaur, Ozan Kumru, Lauren Carter, Jingyou Yu, Aiquan Chang, Katherine McMahan, Thomas Courant, Celia Lebas, Ashley A. Lemnios, Kristen A. Rodrigues, Murillo Silva, Ryan S. Johnston, Christopher A. Naranjo, Mary Kate Tracey, Joseph R. Brady, Charles A. Whittaker, Dongsoo Yun, Swagata Kar, Maciel Porto, Megan Lok, Hanne Andersen, Mark G. Lewis, Kerry R. Love, Danielle L. Camp, Judith Maxwell Silverman, Harry Kleanthous, Sangeeta B. Joshi, David B. Volkin, Patrice M. Dubois, Nicolas Collin, Neil P. King, Dan H. Barouch, Darrell J. Irvine, J. Christopher Love

**Affiliations:** Department of Chemical Engineering, Massachusetts Institute of Technology, Cambridge, Massachusetts 02139, USA; The Koch Institute for Integrative Cancer Research, Massachusetts Institute of Technology, Cambridge, Massachusetts 02139, USA; Department of Biological Engineering, Massachusetts Institute of Technology, Cambridge, Massachusetts 02139, USA; Ragon Institute of MGH, MIT, Harvard, Cambridge, MA 02139, USA; Center for Virology and Vaccine Research, Beth Israel Deaconess Medical Center, Harvard Medical School, Boston, MA, USA; Department of Pharmaceutical Chemistry, Vaccine Analytics and Formulation Center, University of Kansas, Lawrence, Kansas, 66047, United States; Department of Biochemistry, University of Washington, Seattle, WA 98195, USA; Institute for Protein Design, University of Washington, Seattle, WA 98195, USA; Harvard Medical School, Boston, MA 02115, USA; Vaccine Formulation Institute, 1228 Plan-Les-Ouates, Geneva, Switzerland; Harvard-MIT Health Sciences and Technology, Institute for Medical Engineering and Science, Massachusetts Institute of Technology, Cambridge, MA 02139, USA; Bioqual, Inc., Rockville, MD 20850, USA; Bill & Melinda Gates Medical Research Institute, Cambridge, MA 02139, USA; Bill&Melinda Gates Foundation, Seattle, WA 98109, USA; Massachusetts Consortium on Pathogen Readiness, Boston, MA 02115, USA; Howard Hughes Medical Institute, Chevy Chase, MD 20815, USA

## Abstract

Global containment of COVID-19 still requires accessible and affordable vaccines for low- and middle-income countries (LMICs).^1^ Recently approved vaccines provide needed interventions, albeit at prices that may limit their global access.^2^ Subunit vaccines based on recombinant proteins are suited for large-volume microbial manufacturing to yield billions of doses annually, minimizing their manufacturing costs.^3^ These types of vaccines are well-established, proven interventions with multiple safe and efficacious commercial examples.^4–6^ Many vaccine candidates of this type for SARS-CoV-2 rely on sequences containing the receptor-binding domain (RBD), which mediates viral entry to cells via ACE2.^7,8^ Here we report an engineered sequence variant of RBD that exhibits high-yield manufacturability, high-affinity binding to ACE2, and enhanced immunogenicity after a single dose in mice compared to the Wuhan-Hu-1 variant used in current vaccines. Antibodies raised against the engineered protein exhibited heterotypic binding to the RBD from two recently reported SARS-CoV-2 variants of concern (501Y.V1/V2). Presentation of the engineered RBD on a designed virus-like particle (VLP) also reduced weight loss in hamsters upon viral challenge.

---

Vaccines using mRNA have established the efficacy of vaccines for SARS-CoV-2 based on full-length trimeric Spike (S) protein.^9,10^ Recombinant S protein produced in mammalian or insect cells has also shown immunogenicity and efficacy in non-human primates.^11^ For large-volume, low-cost production, however, protein-based vaccines incorporating the receptor binding domain (RBD) subunit remain an important alternative.^12^ Antibodies to RBD account for most of the neutralizing activity elicited in natural infections, and several potent monoclonal antibodies have been discovered from convalescent patients.^13,14^ This domain contains the Receptor-Binding Motif (RBM) that mediates viral entry through the receptor ACE2.^15^ A His-tagged SARS-CoV-2 RBD construct based on SARS-CoV-2 Wuhan-Hu-1 and produced in insect cells has elicited neutralizing antibodies in mice and protective immunity in non-human primates.^16^ Similar tagged constructs have also been adapted for production in yeast like *K. phaffii*,^17,18^ establishing the RBD domain as a prominent candidate for large-volume manufacturing of COVID-19 vaccines.

Despite its significance for low-cost vaccine candidates, recombinant RBD based on the original SARS-CoV-2 clade 19A sequence has shown limited immunogenicity to date. Reported candidates would require as many as three doses or large doses to elicit strong neutralizing antibody responses in mice when formulated with adjuvants.^16,18^ Increasing the number of doses or amounts required could limit its benefits for affordable and accessible vaccines. An engineered design for the RBD, therefore, could enhance the potency of many subunit-based vaccine candidates using this domain.

We reasoned that an improved RBD variant for vaccine candidates should exhibit both improved quality attributes relevant for manufacturing (titers, aggregation) and immunogenicity relative to the Wuhan-Hu-1 sequence used in current vaccines. We further sought to develop a variant amenable to production in microbial hosts, which can be cultured at very large volumes (up to 50,000+ liters) and low costs. Based on previous reports for similar constructs from SARS-CoV-1 and MERS-CoV with demonstrated immunogenicity,^12,19,20^ we first chose to evaluate the production of a tagless 201 amino-acid sequence within the RBD (S protein amino acids 332-532) by secretion from yeast (Figure S1A).

We created a two-stage chromatographic method to purify RBD based on its biophysical characteristics and prior experience purifying heterologous proteins with similar molecular weight, isoelectric point, and hydrophobicity.^21–23^ We produced RBD in 200 mL shake flask culture and purified quantities to assess the quality attributes of the protein (Fig. 1A). The resulting protein bound human ACE2-Fc (K_D_ 49 ± 22 nM) and CR3022 (a neutralizing antibody to SARS-CoV-1 with cross-reactivity to SARS-CoV-2) (K_D_ 32 ± 2 nM) (Fig. S1B-C).^24^ The protein displayed high mannose glycoforms at the single canonical position for *N*-linked glycosylation present on the exposed surface distal from the RBM (Fig. S1D). The protein, however, exhibited a strong tendency to form high-molecular weight species (evident in SDS-PAGE and SEC), particularly in fermentations with high air-water interfaces. The titers were also limited (∼12 mg/L), similar to previously described titers and product quality of unoptimized fermentation.^25^ Together, these results suggested production of this domain was feasible, but presented concerns regarding potential yields and consistency for large-volume manufacturing.

**Fig. 1.**
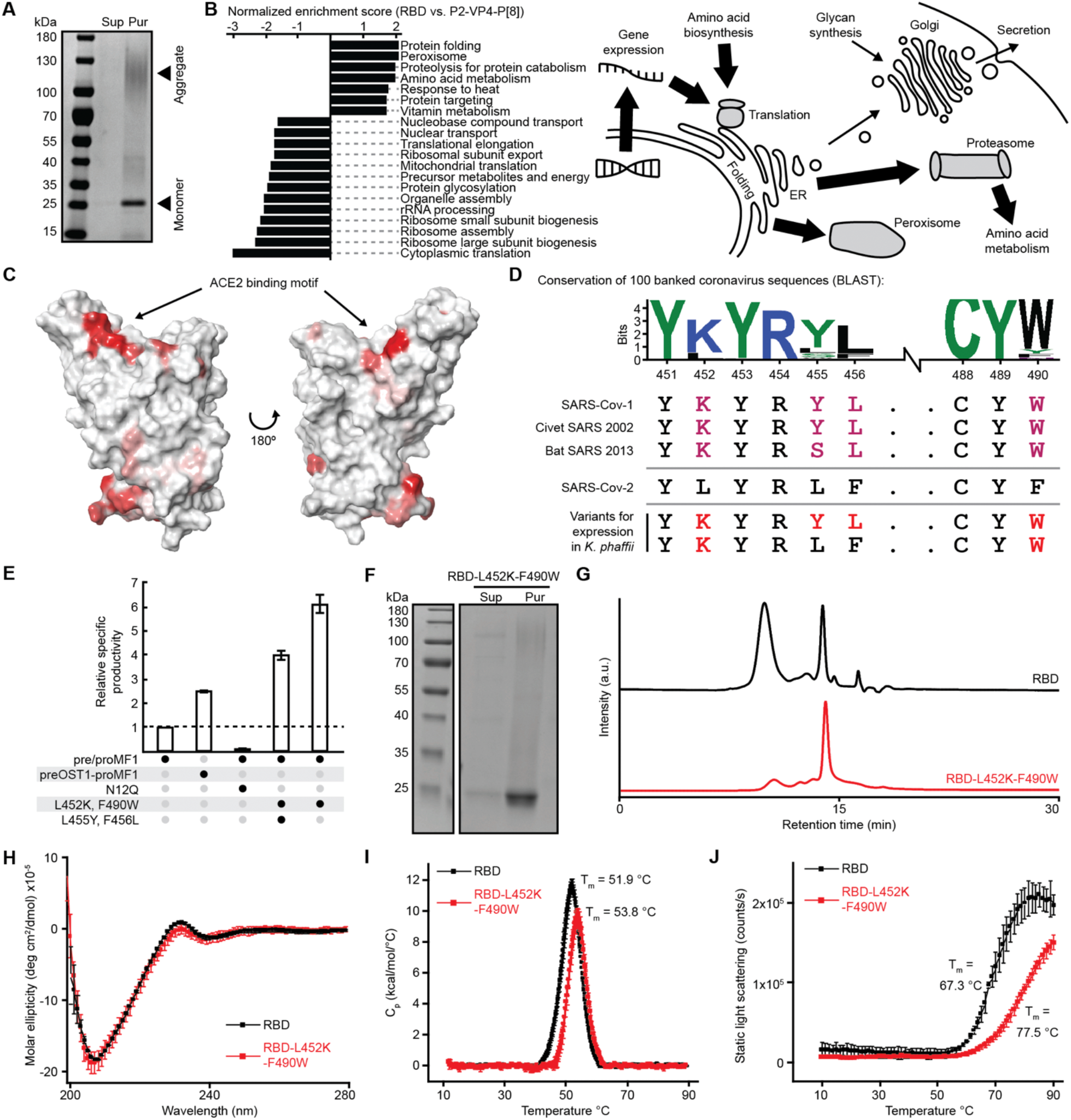
Molecular engineering of the RBD for manufacturability (A) Reduced SDS-PAGE of purified RBD. Sup = cultivation supernatant, Pur = purified protein. (B) Gene set enrichment analysis comparing strains expressing RBD and a rotavirus VP8 fragment (left); schematic model based on pathways for degradation of the RBD in the proteasome and peroxisome, with higher flux of recombinant protein shown with larger arrows (right). (C) Structural rendering of RBD (predicted hydrophobic patches are red). (D) Sequence logo of predicted ACE2 binding motif hydrophobic patch using the top 96 sequences homologous to SARS-CoV-2. Alignment of the ACE2 binding motif to other sarbecoviruses, including selected designs for testing. (E) Bar graph of relative specific productivity for engineered variants of the RBD. preOST1-proMF1 is an alternative signal peptide. Reported values are relative to expression of wild type RBD. (F) Reduced SDS-PAGE of purified RBD-L452K-F490W. (G) Size exclusion chromatography of purified RBD variants. (H) Far-UV circular dichroism at 10° C of purified RBD variants. (I) Differential scanning calorimetry of purified RBD variants. (J) Static light scattering vs. temperature of purified RBD variants.

From these assessments, we reasoned that the qualities of the protein itself may impede its expression and ultimately its attributes that would influence its suitability as an immunogen for subunit-based vaccine candidates. We hypothesized that the tendency of the protein to self-associate may also induce stress on the host cells during the expression and secretion of the protein. Efficient secretion of recombinant protein by yeast requires successful folding and modification of the nascent peptide in the endoplasmic reticulum (ER).^26^ Insoluble or misfolded protein inside the host cells could lead to an unfolded protein response and subsequent degradation of the recombinant product, reducing its production. To further evaluate this relationship, we performed a genome-scale analysis of the yeast by RNA-sequencing and compared the host’s response to the protein to another strain capable of producing a subunit vaccine candidate for rotavirus (P2-VP4-P[8]) of similar size and complexity at commercially-relevant productivities (exceeding 0.5 g/L/d).^23^ Analysis of the differentially regulated pathways revealed differences in gene sets related to protein folding and ER-associated protein degradation pathways. These were upregulated relative to the strain secreting P2-VP4-P[8], implying that the recombinant RBD may be routed from the ER for degradation, reducing yields (Fig. 1B).

Small, conservative changes to a protein sequence can address quality attributes of the protein such as aggregation and also reduce strain on cellular functions to improve titer.^23^ We undertook a similar approach to molecular engineering for SARS-CoV-2 RBD. We inspected the predicted folded structure of the RBD and identified several hydrophobic patches on the surface of this molecule that could promote non-covalent multimerization (Fig. 1C). Spike protein amino acids 452-456 and 488-490 in the RBM had the highest predicted regions of hydrophobicity. To mitigate these hydrophobic patches, we replaced hydrophobic residues with amino acids highly conserved among other sarbecoviruses known to bind ACE2 (Fig. 1D).^27,28^ Lysine residues (as found in other coronaviruses in this region) are generally known to influence adjacent regions in sequences prone to aggregation.^29^ Replacement of only 1-4 amino acids in the RBM *in silico* reduced the AggScore,^30^ a predicted metric of hydrophobicity, of the RBD from 151.26 to 132.46. Based on this analysis, we tested five variants of RBD (Fig. S1E), and found two of these variants (RBD-L452K-F490W and RBD-L452K-L455Y-F456L-F490W) exhibited 4-6 fold increased specific productivity relative to the original strain (Fig. 1E).^31^ We selected the RBD-L452K-F490W to characterize further since it required fewer total changes from the original Wuhan-Hu-1 sequence. We found purified RBD-L452K-F490W exhibited a reduced tendency towards forming high-molecular weight species compared to the original RBD (Fig. 1F-G). We then produced and purified multiple milligrams of each antigen using our InSCyT manufacturing systems for automated, end-to-end production (Fig. S1F-G).^21^ RBD-L452K-F490W exhibited similar secondary structure to the original wildtype sequence (Fig. 1H). The modified sequence manifested a higher melting temperature compared to the original RBD (Fig. 1I), and static light scattering as a function of temperature revealed that thermally induced aggregation of RBD-L452K-F490W was shifted nearly 10°C higher than unmodified RBD (Fig. 1J).

Finally, we reassessed differences in gene expression between strains expressing RBD and RBD-L452K-F490W (Fig. S1H). Contrasting the results for the strain producing the original RBD sequence, the strain expressing RBD-L452K-F490W did not upregulate gene sets related to protein folding and ER-associated protein degradation relative to the strain expressing P2-VP4-P[8], suggesting that the L452K-F490W mutations may alleviate this source of cellular stress. These transcriptomic and biophysical data together suggest the targeted changes to reduce the hydrophobicity of residues within the RBM reduced the propensity for aggregation, enhanced the thermostability of the protein, and improved expression—these traits are all important for large-volume production as well as development of a formulated product with reduced thermal requirements for storage.

The L452K and F490W mutations were selected from conserved substitutions identified from other sarbecoviruses and improved the quality attributes of the RBD, but these changes could alter the antigenicity and immunogenicity of the molecule. Several identified neutralizing antibodies from patients recognize epitopes around the RBM, and many bind near L452.^32^ We measured the affinities of the RBD variant to both human ACE2-Fc and CR3022, and surprisingly, the RBD-L452K-F490W exhibited higher binding affinity to both molecules (K_D_ = 7 ± 1 and 7 ± 1 nM) (Fig. 2A). These data confirmed the engineered RBD variant retains its antigenicity relative to the Wuhan-Hu-1 sequence.

**Fig. 2.**
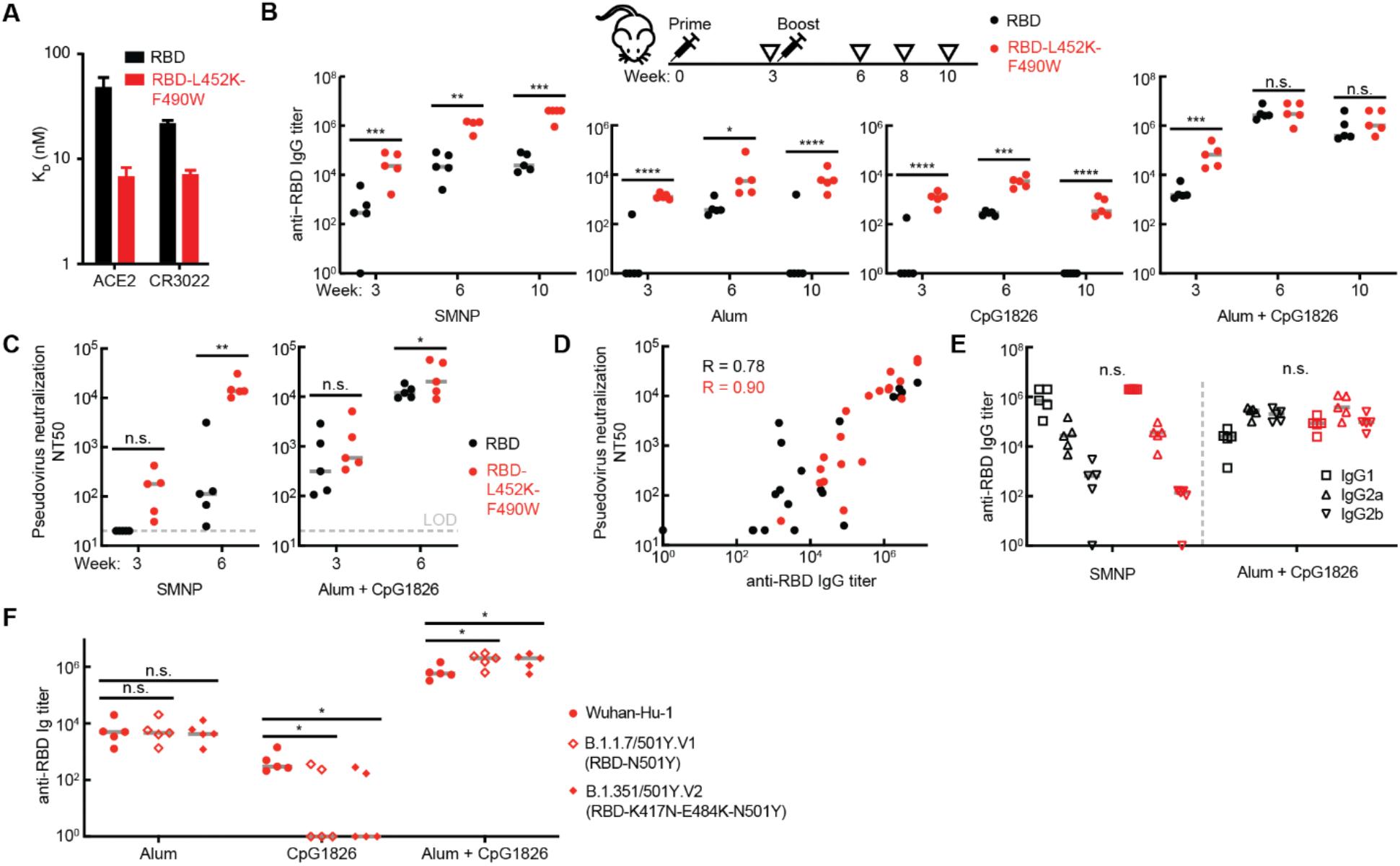
Immunogenicity and antigenicity of wild type and engineered RBD (A) Binding of purified RBD variants to human ACE2-IgG fusion protein and CR3022 neutralizing antibody by biolayer interferometry. (B) Titer of RBD-specific IgG in mouse sera by ELISA. Gray lines represent median values. (C) Titer of neutralizing antibody in mouse sera from SARS-CoV-2 pseudovirus neutralization assay. (D) Correlation of anti-RBD IgG ELISA and pseudovirus neutralization from mouse sera with saponin and alum + CpG adjuvants. (E) Titer of RBD-specific IgG1, IgG2a, and IgG2b antibodies in week 8 mouse sera by ELISA. (F) Titer of RBD-specific IgG in week 8 mouse sera from mice inoculated with RBD-L452K-F490W, evaluated for binding against RBD proteins with mutations from circulating strains of SARS-CoV-2. LOD = limit of detection. Gray lines represent median values. Significance was determined by t-test, with Holm-Sidak correction. *p<0.1, **p<0.01, ***p<0.001, ****p<0.0001

To assess the immunogenicity of our engineered variant, we subcutaneously immunized mice with the unmodified RBD (332-532) or RBD-L452K-F490W formulated with different adjuvants (a saponin-based adjuvant SMNP,^33^ aluminum hydroxide referred to as alum, CpG1826, or a mixture of alum and CpG1826). All animals immunized with RBD-L452K-F490W seroconverted after a single dose, exhibiting robust anti-RBD IgG titers that remained consistently elevated over seven weeks post-boost, regardless of adjuvant (Fig. 2B). In contrast, anti-RBD IgG responses in animals immunized with unmodified Wuhan-Hu-1 RBD were significantly less robust and less durable; serum titers from mice receiving the wildtype sequence with either alum or CpG alone declined to basal levels over time. Furthermore, immunization with RBD-L452K-F490W, both in combination with SMNP and the mixture of alum+CpG adjuvants, elicited pseudovirus neutralizing antibody (NAb) titers after only one dose, with NT50 titers exceeding 10^4^ after a second dose (Fig. 2C-D). These NAb levels were significantly greater than those elicited by the WT sequence both post-prime (3 weeks, SMNP) and post-boost (6 weeks, SMNP and alum+CpG). Of note, SMNP and alum+CpG adjuvant groups elicited high levels of anti-RBD IgG across a distribution of isotypes, including isotypes associated with Th1 (IgG2a and IgG2b) and Th2 (IgG1) responses, suggesting a balanced Th1/Th2 phenotype (Fig. 2D). In contrast, alum alone promoted an IgG1-dominant response, consistent with a Th2 bias (Fig. S2A). Other adjuvants, including MF59, Matrix-M, and aluminum salts have previously been shown to promote functional neutralizing responses for SARS-CoV-1 and MERS.^34^ The RBD-L452K-F490W immunogen also elicited seroconversion in mice similar to full-length S protein when used in combination with oil-in-water emulsion or liposome-based adjuvants (Fig. S2B). Together, these results indicate the engineered variant exhibits enhanced immunogenicity superior to the Wuhan-Hu-1 RBD sequence and could be formulated with several potential adjuvants of commercial relevance.

Antibody responses raised by different antigen-adjuvant combinations can exhibit variable binding and efficacy against naturally occurring variants of SARS-CoV-2.^35^ We tested the binding of antibodies raised against RBD-L452K-F490W to RBD molecules with mutations found in two recently reported SARS-CoV-2 variants of concern, 501Y.V1 and 501Y.V2 (Fig. 2F). Antibodies raised with alum or alum + CpG adjuvants exhibited comparable or slightly improved binding to variant RBDs, while antibodies raised with only CpG adjuvant did not retain binding. These results suggest that immune responses elicited by RBD-L452K-F490W may protect against SARS-CoV-2 variants with the N501Y spike protein mutation.

Multimeric display of subunit antigens like RBD on nanoparticle-based scaffolds provides a promising approach to enhance immunogenicity further and to reduce the amount of protein required for individual doses of a vaccine or the number of doses required.^36,37^ Both attributes could facilitate broader global coverage for COVID-19 vaccines. We further modified the engineered RBD-L452K-F490W to include a peptide motif for covalently linking the antigen to a virus-like particle (VLP) via a transpeptidation reaction and produced the antigen similarly to the unmodified version (Fig. 3A,B; Fig. S3A).^38,39^ We conjugated the engineered antigen onto a designed self-assembling nanoparticle (i3-01) produced in bacteria.^40^ The resulting particles had ∼85% occupancy of the 60 available sites for antigenic display on each VLP (Fig. 3A-B, Fig. S3C). We confirmed that VLPs were correctly assembled by electron microscopy and size exclusion chromatography before and after conjugation (Fig. 3C, Fig. S3D). We immunized mice with these constructs with doses containing 5 µg of RBD down to 0.06 µg, with alum and CpG adjuvants. All doses induced seroconversion with a strong correlation evident between the anti-RBD Ig response and neutralizing titers (Fig. 3D-E, Fig. S3D). We then immunized golden Syrian hamsters with the RBD-decorated VLPs with either alum or alum and CpG1018—a commercial GMP-grade adjuvant—with a prime and a boost after 3 weeks. Following the boost, we challenged the hamsters with SARS-CoV-2 and monitored for body weight change and viral titer post challenge. Animals that received the RBD-VLP with alum+CpG recovered in weight faster than the control group (p=0.04, day 6 post challenge) (Fig. 3F, Fig. S3E-F). Across all vaccinated animals, body weight change correlated with the measured titer of neutralization antibody from sera (Fig. S3G). These two studies demonstrate one potential presentation of the RBD-L452K-F490W as a vaccine antigen on a nanoparticle, and its efficacy reducing the effects of SARS-CoV-2 in the hamster model.

**Fig. 3.**
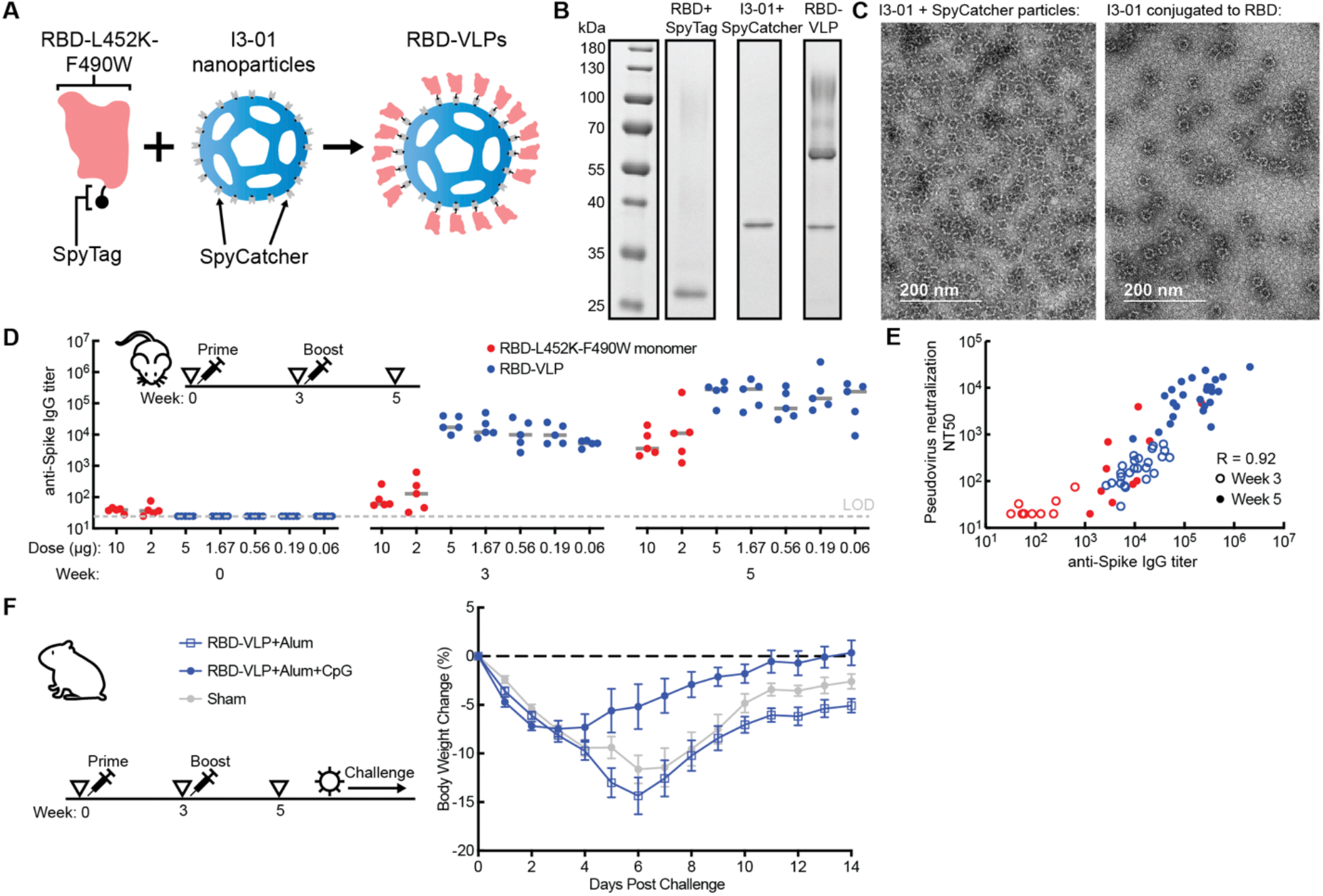
Immunogenicity and antigenicity of engineered RBD nanoparticles in mice and hamsters (A) Schematic of nanoparticle assembly using SpyTag and SpyCatcher. (B) Reduced SDS-PAGE of nanoparticle components. (C) Negative stain electron microscopy of SpyCatcher-12GS-I3-01 nanoparticles before (left) and after (right) conjugation to RBD-L452K-F490W-GGDGGDGGDGG-SpyTag. (D) Titer of spike protein-specific IgG in mouse sera by ELISA. (E) Spearman correlation of anti-S protein IgG ELISA and pseudovirus neutralization from mouse sera. (F) Mean percent body weight change of hamsters in each group after challenge with SARS-CoV-2. Gray bars represent median values.

In conclusion, we have demonstrated an engineered variant of SARS-CoV-2 RBD that shows improved immunogenicity compared to the original Wuhan-Hu-1 sequence for RBD used in current vaccines and is compatible with multiple commercially-relevant adjuvants. This design also exhibits improved biomolecular attributes that make it well-suited for further development for large-volume manufacturing of low-cost vaccine candidates. Improving the designs of vaccines for COVID-19 will remain critical as new variants like 501Y.V1/V2 continue to emerge with mutations in the RBD domain. Such adaptations by the virus could reduce the effectiveness of interventions like monoclonal antibodies and current vaccines based on the original Wuhan-Hu-1 strain.^41^ Antibodies raised against the engineered RBD reported here exhibit heterotypic binding to both 501Y.V1 and V2. Interestingly, these viral variants also exhibit enhanced binding to ACE2, similar to the design here,^42^ and genomic sequences for reported strains containing mutations at E484 also show co-occurrences with changes at F490.^43^ The ACE2 RBM remains a critical epitope for neutralization of emerging variants.^44^ Another recent variant (B.1.429/CAL.20C) that also shows escape from known neutralizing antibodies for SARS-CoV-2 contains a strikingly similar change in the first position identified in our engineered design (L452R).^45^ The modifications to RBD we demonstrated here are limited to the ACE2 RBM, and, therefore, are compatible in principle with any vaccine based on presentation of the S protein or RBD, and could be combined with other specific mutations identified from naturally occurring variants. The increased immunogenicity of the design presented here may afford further insights to improve the breadth of protection afforded by SARS-CoV-2 vaccine candidates, and ultimately affordable and accessible vaccines.

---

**Fig. S1.**
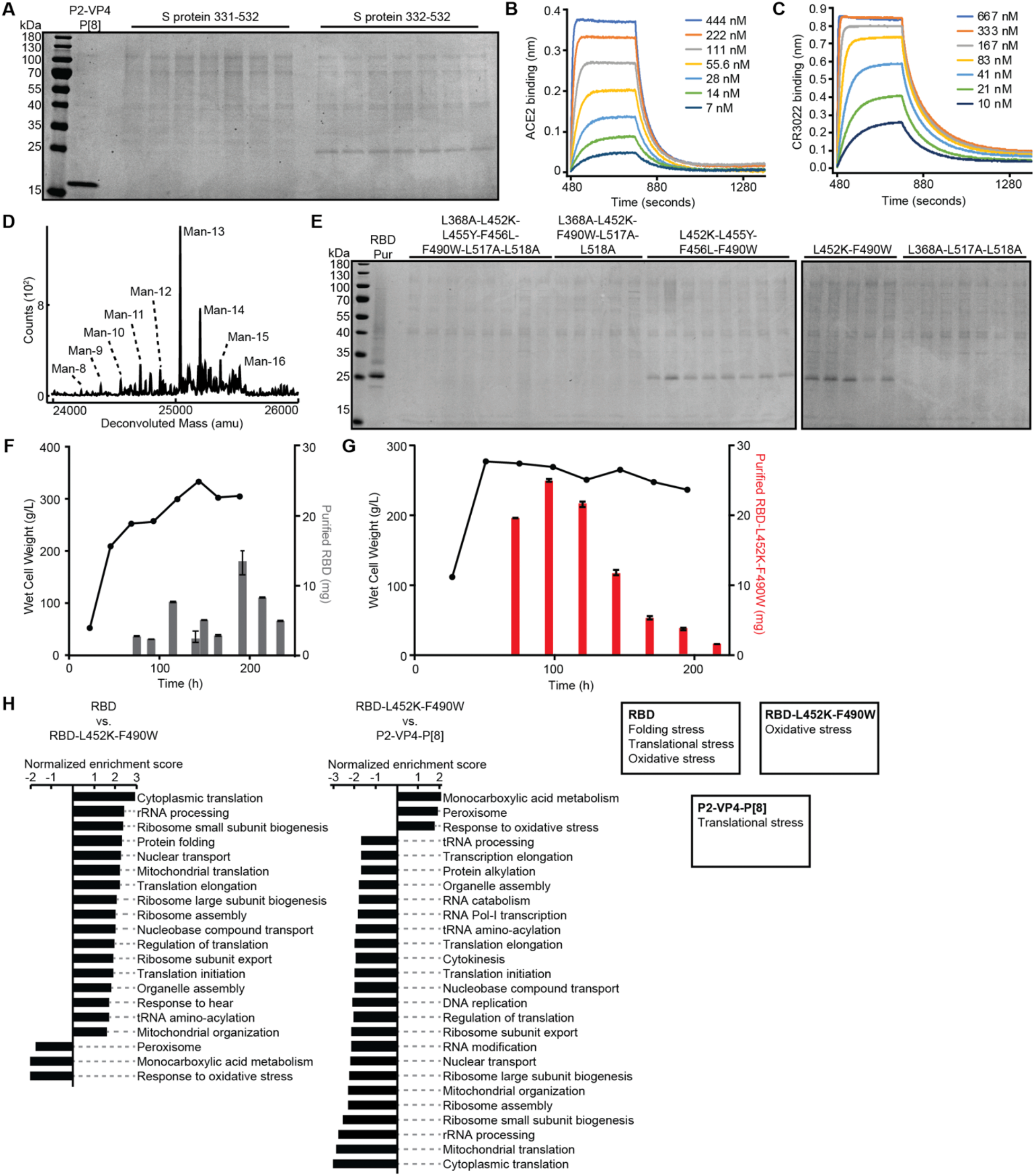
Initial expression and manufacturing of RBD variants (A) Reduced SDS-PAGE of cultivation supernatants of initial RBD variants. Each lane represents a unique clone after transformation. (B,C) Binding of RBD to human ACE2-IgG fusion protein (B) and CR3022 neutralizing antibody (C) by biolayer interferometry. (D) Mass spectrum of purified RBD with labeled specific glycan peaks. Man = mannose. (E) Reduced SDS-PAGE of cultivation supernatants of RBD variants. (F-G) Yields for production of RBD (F) and RBD-L452K-F490W (G). Wet cell weight and purified pools of RBD variants are shown. (H) Gene set enrichment analysis comparing strains expressing RBD, RBD-L452K-F490W, and a rotavirus VP8 fragment (left); summary of upregulated cellular processes in each strain (right).

**Fig. S2.**
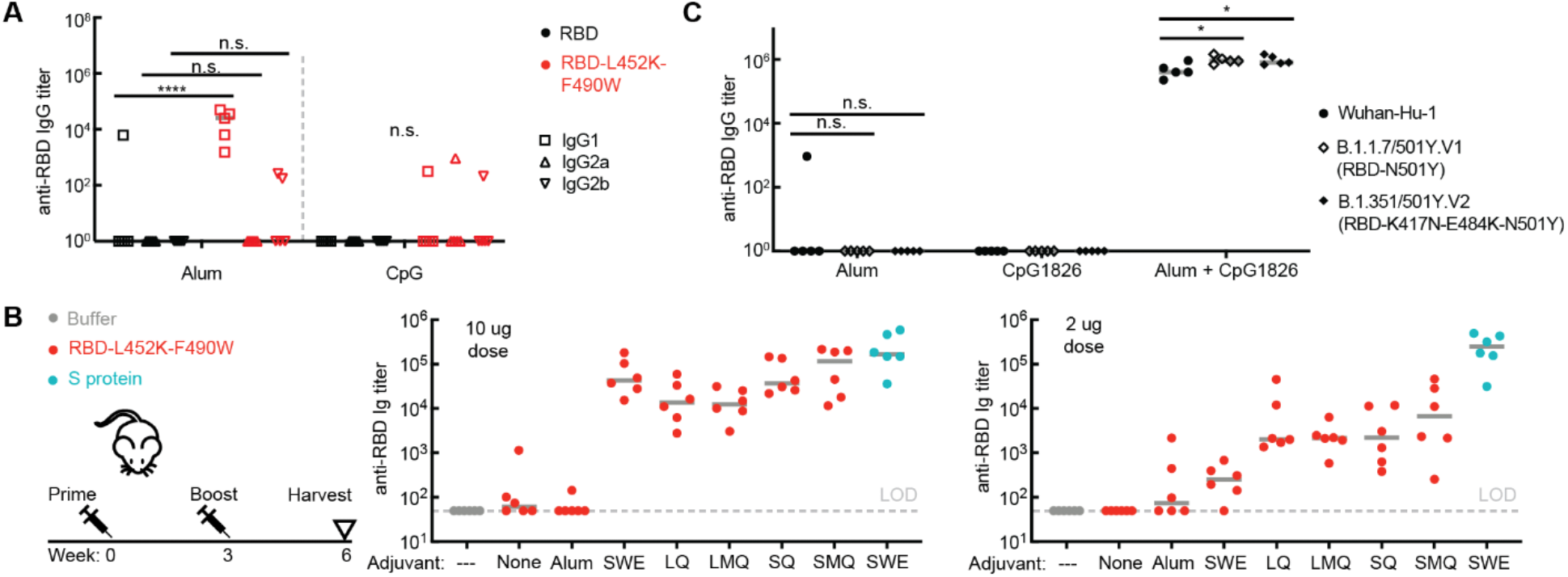
Immunogenicity of engineered RBD formulated with adjuvants (A) Titer of RBD-specific IgG1, IgG2a, and IgG2b antibodies in mouse sera by ELISA. (B) Titer of RBD-specific total Ig in mouse sera by ELISA. RBD-L452K-F490W was administered in 2 µg or 10 µg doses at weeks 0 and 3 with different adjuvants. (C) Titer of RBD-specific IgG in week 8 mouse sera from mice inoculated with RBD, evaluated for binding against RBD proteins with mutations from circulating strains of SARS-CoV-2. Gray lines represent median values. Significance was determined by t-test, with Holm-Sidak correction. *p<0.1.

**Figure S3.**
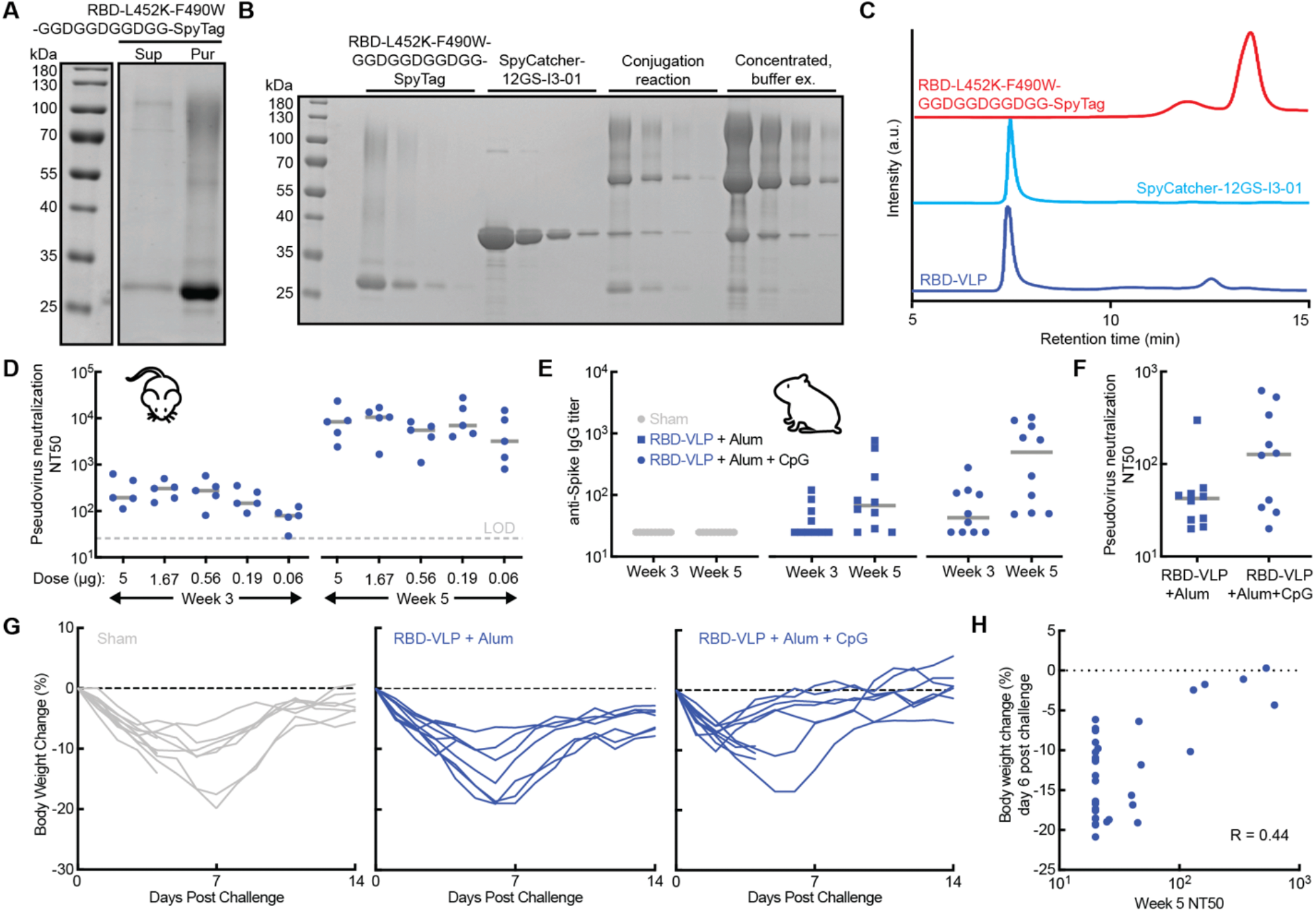
Production and characterization of RBD-VLP nanoparticles (A) Reduced SDS-PAGE of RBD-L452K-F490W-GGDGGDGGDGG-SpyTag purification. Sup = cultivation supernatants, Pur = purified protein. (B) Reduced SDS-PAGE of purified RBD-L452K-F490W-GGDGGDGGDGG-SpyTag, SpyCatcher-12GS-I3-01 (produced in *E. coli*), nanoparticles after the conjugation reaction, and nanoparticles after buffer exchange and filter concentration. Gel replicates are serial 3x dilutions of each sample. (C) Size exclusion chromatography of nanoparticles before and after the conjugation reaction. (D) Titer of neutralizing antibody in mouse sera from SARS-CoV-2 pseudovirus neutralization assay for the RBD-VLP dosing study. (E) Titer of spike protein-specific IgG in hamster sera by ELISA. Doses were administered at weeks 0 and 3. (F) Titer of neutralizing antibody in hamster sera from SARS-CoV-2 pseudovirus neutralization assay. (G) Change in body weight of individual hamsters after challenge with SARS-CoV-2. Three animals in each group underwent scheduled necropsies at day 4. (H) Correlation of percentage weight change at day 6 post challenge with neutralizing antibody titer in sera sampled in week 5. Gray lines represent median values. LOD = limit of detection.

## Acknowledgements

We thank Prof. Ragahavan Varadarajan of IISc for kindly providing the RBD sequence for SARS-CoV-2 Wuhan-Hu-1. We thank Laurent Pessaint, Dr. Adrienne Winn, Kamil Radzyminski, Andrew Faudree, Brittany Spence, Melissa Hamilton, and Natalie Figueroa Jones for advice and assistance with hamster studies. We thank Sumi Biswas of SpyBiotech for helpful discussions on SpyTag/SpyCatcher conjugations. The authors thank the Koch Institute’s Robert A. Swanson (1969) Biotechnology Center for technical support. The following reagent was deposited by the Centers for Disease Control and Prevention and obtained through BEI Resources, NIAID, NIH: SARS-Related Coronavirus 2, Isolate USA-WA1/2020, NR-53780. This work was funded by the Bill and Melinda Gates Foundation (Investment IDs INV-002740 and INV-006131). This study was also supported in part by the Koch Institute Support (core) Grant P30-CA14051 from the National Cancer Institute. L.H.T. is an NIH T32 Postdoctoral Fellow supported by the Multidisciplinary AIDS Training Program (Grant # T32 AI007387). The content is solely the responsibility of the authors and does not necessarily represent the official views of the NCI, the NIH, Gates MRI, or the Bill and Melinda Gates Foundation.

## Author contributions

N.C.D., S.R.A., and J.C.L. conceived and planned experiments. N.C.D. and R.S.J. generated and characterized yeast strains. N.C.D., A.M.B., J.R.B., and C.A.W. performed RNA sequencing. A.M.B. and M.K.T. conducted bioreactor experiments. S.R.A. and A.M.B. performed HPLC assays. S.R.A. and L.E.C. designed and performed protein purifications. N.C.D. and C.A.N. performed mass spectrometry. K.K. and O. K. performed biophysical characterization. L.C. and N.P.K. produced and purified nanoparticle proteins. D.S.Y. performed electron microscopy. N.C.D. and S.R.A. formulated samples for animal studies. B.L.H., A.A.L., and K.A.R. designed and performed soluble RBD mouse studies and RBD ELISA assays. M.S. produced the SMNP adjuvant. T.C., C.L., P.M.D., and N.C. designed and performed adjuvant mouse studies. L.H.T., K.M., and A.C. designed and performed RBD-VLP mouse studies. L.H.T., A.C., S.K., M.P., M.L., H.A., and M.G.L. designed and performed RBD-VLP hamster studies. J.Y., A.C., and L.H.T. performed pseudovirus neutralization assays. J.M.S. and D.L.C. coordinated and managed resource allocations and planning for experimental studies. N.C.D., S.R.A., K.R.L., and J.C.L. wrote the manuscript. J.C.L., D.J.I., D.B.V., D.H.B., N.P.K., P.M.D., N.C., S.B.J., K.R.L. J.M.S. and H.K. designed the experimental strategy and reviewed analyses of data. All authors reviewed the manuscript.

## Materials and Methods

### Strains

All strains were derived from wild-type *Komagataella phaffii* (NRRL Y-11430), in a modified base strain (RCR2_D196E, RVB1_K8E) described previously.^1^ Genes containing RBD variants were codon optimized, synthesized (Integrated DNA Technologies), and cloned into a custom vector. *K. phaffii* strains were transformed as described previously.^2^

### Cultivations

Strains for initial characterization and titer measurement were grown in 3 mL culture in 24-well deep well plates (25°C, 600 rpm), and strains for protein purification were grown in 200 mL culture in 1 L shake flasks (25°C, 250 rpm). Cells were cultivated in complex media (potassium phosphate buffer pH 6.5, 1.34% nitrogen base w/o amino acids, 1% yeast extract, 2% peptone). Cells were inoculated at 0.1 OD600, outgrown for 24 h with 4% glycerol feed, pelleted, and resuspended in fresh media with 3% methanol to induce recombinant gene expression. Supernatant samples were collected after 24 h of production, filtered, and analyzed. InSCyT bioreactors were operated as described previously.^3^

### Transcriptome analysis

Cell were harvested after 18 h of production at 3 mL plate scale. RNA was extracted and purified according to the Qiagen RNeasy kit (cat #74104) and RNA quality was analyzed to ensure RNA Quality Number >6.5. RNA libraries were prepared using the 3’DGE method and sequenced on an Illumina Nextseq to generate paired reads of 20 (read 1) and 72 bp (read 2). Sequenced mRNA transcripts were demultiplexed using sample barcodes and PCR duplicates were removed by selecting one sequence read per Unique Molecular Identifier (UMI) using a custom python script. Transcripts were quantified with Salmon version 1.1.0^4^ and selective alignment using a target consisting of the *K. phaffii* transcripts, the RBD-N1del, P[8] and selectable marker transgene sequences and the *K. Phaffii* genome as a selective alignment decoy. Expression values were summarized with tximport version 1.12.3^5^ and edgeR version 3.26.8.^6,7^ Expression was visualized using *log*_*2*_*(Counts per Million + 1)* values. Gene set enrichment analysis (GSEA) was performed with GSEA 4.1.0 using Wald statistics calculated by DESeq2^8^ and gene sets from yeast GO Slim.^9^

### Protein purification

Protein purification for non-clinical studies and end-to-end manufacturing was carried out on the purification module of the InSCyT system as described previously.^3^ All columns were equilibrated in the appropriate buffer prior to each run. Product-containing supernatant was adjusted to pH 4.5 using 100mM citric acid. The adjusted supernatant was loaded into a pre-packed CMM HyperCel column (5-mL) (Pall Corporation, Port Washington, NY), re-equilibrated with 20 mM sodium citrate pH 5.0, washed with 20 mM sodium phosphate pH 5.8, and eluted with 20 mM sodium phosphate pH 8.0, 150 mM NaCl. Eluate from column 1 above 15 mAU was flowed through a 1-mL pre-packed HyperCel STAR AX column (Pall Corporation, Port Washington, NY). Flow-through from column 2 above 15 mAU was collected.

### Analytical assays for protein characterization

Purified protein concentrations were determined by absorbance at A280 nm. SDS-PAGE was carried out as described previously.^3^ Supernatant titers were measured by reverse phase liquid chromatography, and normalized by cell density, measured by OD600.

### Biolayer interferometry

Biolayer interferometry was performed using the Octet Red96 with Protein A (ProA) biosensors (Sartorius ForteBio, Fremont, CA), which were hydrated for 15 min in kinetics buffer prior to each run. Kinetics buffer comprising 1X PBS pH 7.2, 0.5% BSA, and 0.05% Tween 20 was used for all dilutions, baseline, and disassociation steps. CR3022 and ACE2-Fc were used in the assay at concentrations of 2 and 10 µg/mL, respectively. Samples were loaded in a 96-well black microplate (Greiner Bio-One, Monroe, NC) at starting concentrations of 15 and 10 µg/mL, respectively. Seven 1:1 serial dilutions and a reference well of kinetics buffer were analyzed for each sample. Association and dissociation were measured at 1000 rpm for 300 and 600 sec, respectively. Binding affinity was calculated using the Octet Data Analysis software v10.0 (Pall ForteBio), using reference subtraction, baseline alignment, inter-step correction, Savitzky-Golay filtering, and a global 1:1 binding model.

### Size exclusion chromatography

Size exclusion high performance liquid chromatography (HPLC) analysis was performed on an Agilent 1260 HPLC system controlled using OpenLab CDS software (Agilent Technologies, Santa Clara, CA). The analysis was performed using an AdvanceBio SEC column (Agilent Technologies, Santa Clara, CA, 4.6 x 300 mm, 300Å, 2.7µm) and AdvanceBio SEC guard column (Agilent Technologies, Santa Clara, CA, 4.6 x 50 mm, 300Å, 2.7µm). The column was operated at a flow rate of 0.25 mL/minute and ambient temperature. The mobile phase buffer was 150mM sodium phosphate (Sigma-Aldrich, St. Louis, MO), pH 7.0. Total method run time was 30 minutes and sample injection volumes were 10µL. A diode array detector was set for absorbance detection at 214nm. Data analysis was completed using Agilent’s OpenLab CDS Data Analysis.

### Reverse phase chromatography

Reverse phase high performance liquid chromatography (HPLC) analysis was performed on an Agilent 1260 HPLC system controlled using OpenLab CDS software (Agilent Technologies, Santa Clara, CA). Antigen concentration was determined using a PLRP-S column (2.1 x 150 mm, 300Å, 3µm) operated at 0.6 mL/min and 80°C (Agilent Technologies, Santa Clara, CA) on an HPLC equipped with a diode array detector set for absorbance detection at 214nm and. Buffer A was 0.1% (v/v) TFA in water and buffer B was 0.1% (v/v) TFA, 0.5% (v/v) water in ACN. A gradient of 39-43% B was performed over 9 minutes; total method run time was 18 minutes. Sample injection volumes were 50µL. A diode array detector was set for absorbance detection at 214nm. Data analysis was completed using OpenLab CDS Data Analysis (Agilent Technologies, Santa Clara, CA).

### Mass spectrometry

Intact mass analysis was performed on a 6530B Q-TOF LC-MS with a 1290 series HPLC (Agilent Technologies, Santa Clara, CA). Mobile phase A consisted of LC-MS grade water with 0.1% formic acid, and mobile phase B was LC-MS grade acetonitrile with 0.1% formic acid. About 1.0 µg of protein for each sample was injected, bound to a PLRP-S column (2.1mm x 50mm, 5μm, 300Å) (Agilent Technologies), desalted, and subjected to electrospray ionization. The LC gradient comprised 5-95% B over 30 min at a flow rate of 0.4 mL/min. A blank injection between each sample was performed as a wash step, consisting of an LC gradient of 5-95% B over 6 min. The electrospray ionization parameters were: 325°C drying gas temperature, 10 L/min drying gas flow, 30 psig nebulizer, 4000 V Vcap, and 325 V fragmentor voltage. Mass spectra were collected from 500-3200 m/z at a scan rate of 1 spectra/sec. MS spectra were processed using MassHunter Bioconfirm software (v B.10.0, Agilent Technologies) with a deconvolution range of 10-50 kDa, using a mass step of 1 Dalton.

### Far-UV Circular Dichroism (CD)

CD spectroscopy was performed using a Chirascan-plus CD spectrometer (Applied Photophysics Ltd., Leatherhead, UK) equipped with a 6-cuvette position Peltier temperature controller (Quantum Northwest, Liberty Lake, WA) and a high-performance solid-state detector. The lamp (150 W air-cooled Xe arc) housing, monochromator and sample compartment were continuously purged with N_2_ gas. The 10°C CD spectra of RBD samples at 0.2 mg/mL were collected in triplicate in the range of 280-200 nm using quartz cuvettes (1 mm path length) sealed with a Teflon stopper (Starna Cells Inc., Atascadero, CA). Data were subjected to a 3-point Savitzky-Golay smoothing filter using the Chirascan software (Applied Photophysics) and the ellipticity of the buffer was subtracted from all sample measurements.

### Static Light Scattering vs. Temperature

Static light scattering measurements as a function of temperature were made in triplicate using a dual emission PTI QM-40 Spectrofluorometer (Horiba Scientific Northampton, UK) equipped with a 4-position cell holder Peltier temperature control device, a high-power continuous 75 W short-arc Xe lamp (Ushio), and a Hamamatsu R1527 photomultiplier tube. Data were collected using FelixGX software (Horiba Scientific) in 10 mm path length quartz cuvettes. RBD samples at 0.2 mg/mL were examined as a function of temperature (10°C-90°C) using an excitation wavelength of 295 nm. Static light scattering signal at 295 nm was collected at 1.25°C interval with a 2 min equilibration at each temperature. The light scattering signal of the buffer was subtracted from all sample measurements and the light scattering intensity at 295 nm was plotted at a function of temperature.

### Differential Scanning Calorimetry (DSC)

DSC was performed in triplicate using an auto-VP capillary differential scanning calorimeter (MicroCal/GE Health Sciences, Pittsburgh, PA) equipped with Tantalum sample and reference cells pressurized at ∼60 psi with nitrogen. RBD samples at 0.2 mg/mL were loaded in the autosampler tray held at 4°C and scans were completed from 10°C to 90°C using a scan rate of 60°C/h and a pre-scan thermostat of 15 min. Buffer subtraction and concentration normalization were performed using Origin (OriginLab, Northampton, MA). Data analysis was performed using the MicroCal LLC DSC plug-in for the Origin 7.0 software package.

### Immunization of mice for soluble RBD studies

The immunogenicity of RBD-L452K-F490W compared to RBD was evaluated *in vivo* in mice. All procedures were approved by the Massachusetts Institute of Technology Institutional Animal Care and Use Committee (IACUC) following local, state, and federal regulations. Immunization studies were carried out using age-matched 6-8 wk old Balb/cJ female mice purchased from The Jackson Laboratory (strain: 000651). Mice were immunized on day 0 and day 21 with 5µg RBD plus adjuvant: 50µg alum Alhydrogel (Invivogen), 30µg CpG1826 (Invivogen), or 5µg saponin MPLA nanoparticles (SMNP). SMNP was synthesized in-house as previously described,^10^ where dose is reported as the amount of saponin administered. Immunizations were administered via subcutaneous injection in 100µl PBS at the tail base (2 x 50µl bilateral injections, one on each side of the tail base). Blood was collected by cheek or retro-orbital bleed for ELISA antibody analysis on wk 2, 3, 4, and then every 2 weeks thereafter. Serum was isolated from blood using serum separator tubes, centrifuged at 10,000xg for 5min at 4C, then stored at -80C.

### RBD-specific ELISA assays

Anti-RBD IgG was measured in mouse serum by ELISA. To capture serum antibodies from immunized mice, Costar Polystyrene High Binding 96-well plates (Corning) were coated directly with RBD antigen at 2µg/ml in PBS overnight at 4C, then blocked with PBS + 2% BSA for 2 hr at 25C. Mouse sera were diluted in block buffer (PBS + 2% BSA) starting at 1:100 or 1:200 followed by 4X serial dilutions and incubated in plates for 2 hr at 25C, followed by detection with 1:5000 goat anti-mouse IgG-HRP (BioRad) in block buffer for 1 hr. Plates were developed using TMB substrate for 1-20 min and stopped with 2N sulfuric acid. For all titer analyses, samples directly compared across groups were developed for the same amount of time. Cut-off titers are reported as inverse dilutions giving a 0.2 HRP absorbance (A450 – A540).

### Immunization of mice for RBD adjuvant studies

The immunogenicity of RBD-L452K-F490W in combination with various adjuvants was evaluated in 7-8 week-old C57BL/6J female mice (Charles River, strain: 000634) at the Vaccine Formulation Institute (VFI, Switzerland). All animal work was performed in accordance with the Swiss Federal Animal Protection Act. Mice were immunized IM with RBD antigen and adjuvants including aluminum Hydroxide (AlOH, Croda, Denmark), SWE^11^ (squalene-in-water emulsion, Seppic, France), SQ (SWE + QS21 saponin, VFI), SMQ^12^ (squalene-in-water emulsion + synthetic TLR4L and QS21, VFI), LQ^12^ (neutral liposomes + QS21, VFI) or LMQ^12^ (neutral liposome + synthetic TLR4L and QS21 saponin, VFI). The TLR4 agonist was used at a final concentration of 2 µg/dose and the QS21 saponin at 5 µg/dose. The Spike prefusion trimer (produced previously^13^) adjuvanted with SWE was used as a benchmark in this experiment as it has been shown to induce high titers of neutralizing antibodies. All formulations were fully characterized for adjuvant physico-chemical properties and antigen integrity. Blood samples were collected on day 42 and sera tested for RBD specific antibodies in ELISA with the following modifications: Plates were coated with 1.25µg/mL soluble RBD (produced previously^13^) and mouse serum Ig was detected using a goat anti mouse Ig coupled to HRP (Southern Biotech) diluted at 1/6000.

### Production of RBD nanoparticles

I3-01-spycatcher nanoparticles were manufactured in *E. coli* using standard methods. Protein was expressed using standard IPTG methods, using pet29B expression vector and NEB Lemo21 expression strain. Cells were lysed with a Microfluidics M110P microfluidizer, clarified by centrifugation, and captured from soluble lysate by IMAC, using Cytiva IMAC-FF Sepharose resin charged with NiSO4. Protein of interest was eluted from resin using imidazole, concentrated, and polished using Size Exclusion Chromatography on a Cytiva Superose 6 Increase column. Assembly of the nanoparticle was confirmed using Dynamic Light Scattering and nsEM, in conjunction with the observed SEC retention time. Endotoxin was removed using Pierce™ High Capacity Endotoxin Removal Spin Columns (Thermo Scientific) to below 10 EU/mL and quantified using Pierce™ Chromogenic Endotoxin Quant Kit (Thermo Scientific). I3-01-spycatcher and RBD-spytag were conjugated by incubation overnight at 4°C in 20mM sodium phosphate, 150mM NaCl, pH 8 buffer, with a 1:1.5 I3-01-spycatcher:RBD-spytag molar ratio. Excess RBD was removed with a 100 kDa molecular weight cutoff Amicon® Ultra-4 centrifugal filter (Millipore). RBD valency was determined using SDS-PAGE densitometry in ImageJ.

### Negative-stain electron microscopy

Solution with conjugated RBD-VLP nanoparticles (7 µL) was incubated on a 200 meshes copper grid coated with a continuous carbon film for 60 seconds. Excess liquid was removed, and the film was incubated in 10 µL of 2% uranyl acetate. The grid was dried at room temperature and mounted on a JEOL single tilt holder in the TEM column. The specimen was cooled by liquid nitrogen, and imaged on a JEOL 2100 FEG microscope with a minimum dose to avoid sample damage. The microscope was operated at 200 kV with magnification at 10,000-60,000x. Images were recorded on a Gatan 2Kx2K UltraScan CCD camera.

### Immunization of mice for RBD-VLP studies

Balb/cJ female mice, ages 6-8 weeks were purchased from The Jackson Laboratory (strain: 000651). Mice were immunized on day 0 and day 21 with the indicated doses of RBD-VLPs or soluble RBD-L425K-F490W monomer as a control; all groups were adjuvanted with 40µg Alum + 10µg CpG. Immunizations were administered via bilateral intra-muscular injection. Peripheral blood was collected via the submandibular route at day 0, day 21, and day 35, and serum was isolated for immunologic assays. Studies were conducted in compliance with all relevant local, state and federal regulations and were approved by the Beth Israel Deaconess Medical Center Institutional Animal Care and Use Committee.

### Spike protein-specific ELISA assays

Nunc Immuno MaxiSorp 96-well plates (Thermo Scientific) were coated with SARS-CoV-2 Spike protein (Sino Biological) at 1µg/mL in PBS and incubated overnight at 4C. After incubation, plates were washed once with wash buffer (0.05% TWEEN-20 in 1X PBS), then blocked with casein for 2-3 hours at 25C. Mouse or hamster sera were diluted in casein block buffer starting at 1:25 followed by 3X serial dilutions and incubated in plates for 1 hr at 25C. Plates were then washed three times, and incubated with either rabbit anti-mouse IgG-HRP (Jackson Immuno) or goat anti-hamster IgG(H+L)-HRP (SouthernBiotech) diluted in casein for mouse or hamster samples, respectively. After 1 hour incubation at 25C, plates were wsaahed three times, then developed using TMB substrate (SeraCare). Development was halted using stop solution (SeraCare). For each sample, ELISA endpoint titer was calculated in Graphpad Prism software, using a four-parameter logistic curve fit to calculate the reciprocal serum dilution that yields an absorbance value (450nm) of 0.2.

### Pseudovirus neutralization assays

The SARS-CoV-2 pseudoviruses expressing a luciferase reporter gene were generated in an approach similar to as described previously.^14–16^ Briefly, the packaging plasmid psPAX2 (AIDS Resource and Reagent Program), luciferase reporter plasmid pLenti-CMV Puro-Luc (Addgene), and spike protein expressing pcDNA3.1-SARS CoV-2 SΔCT were co-transfected into HEK293T cells by lipofectamine 2000 (ThermoFisher). The supernatants containing the pseudotype viruses were collected 48 h post-transfection, which were purified by centrifugation and filtration with 0.45 µm filter. To determine the neutralization activity of the plasma or serum samples from animals, HEK293T-hACE2 cells were seeded in 96-well tissue culture plates at a density of 1.75 × 10^4^ cells/well overnight. Three-fold serial dilutions of heat inactivated plasma samples were prepared and mixed with 50 µL of pseudovirus. The mixture was incubated at 37°C for 1 h before adding to HEK293T-hACE2 cells. 48 h after infection, cells were lysed in Steady-Glo Luciferase Assay (Promega) according to the manufacturer’s instructions. SARS-CoV-2 neutralization titers were defined as the sample dilution at which a 50% reduction in relative light unit (RLU) was observed relative to the average of the virus control wells.

### Immunization of hamsters

Male and female Syrian golden hamsters (Envigo), 7-8 weeks of age, were randomly distributed into three groups. Hamsters were immunized via intramuscular injection with a prime at week 0, followed by a boost at week 3 of i) RBD-J-ST-i3 VLPs containing 2µg of RBD protein + 40µg of Alum, ii) RBD-J-ST-i3 VLPs containing 2µg of RBD protein + 40µg of Alum + 100µg of CpG, or iii) sham. Peripheral blood was drawn via the retro-orbital route at baseline (week 0), week 3, and week 5 to collect serum for immunologic assays. At week 5, hamsters were challenged with 1.99 × 10^4^ TCID50 SARS-CoV-2, derived from USA-WA1/2020 (NR-53780, BEI Resources). Challenge virus was administered in 100µL of total volume by the intranasal route (50µL in each nare). After challenge, body weights of hamsters were monitored daily. Studies were conducted in compliance with all relevant local, state and federal regulations and were approved by the Bioqual Institutional Animal Care and Use Committee.

